# Steering fatty acid composition of yeast microbial oil via genetic modification and bioprocess adjustment

**DOI:** 10.1101/2025.03.14.643280

**Authors:** Zeynep Efsun Duman-Özdamar, Edoardo Saccenti, Vitor A.P. Martins dos Santos, Jeroen Hugenholtz, Maria Suarez-Diez

## Abstract

The increasing demand for palm oil has drastic effects on the ecosystem as its production is not sustainable. To that end, developing a sustainable alternative to fatty acids and oils is urgent and of utmost interest. Oils produced by oleaginous yeasts present a promising solution, particularly because the fatty acid profile of the oil produced by these yeasts is comparable to that of plant-based oils and fats. The fatty acid composition of the oil determines its physiological properties, thereby determining its potential applications. Accordingly, the production of microbial oil with an optimal composition profile for a specific application is of great importance. In this study, we evaluated the variation that occurred in fatty acid composition due to different cultivation parameters (temperature, C/N ratio, carbon, and nitrogen sources) and applied genetic modifications to improve the lipid accumulation of *Cutaneotrichosporon oleaginosus* and *Yarrowia lipolytica*. We showed that specific fatty acid profiles associated with a particular application can be obtained by carefully selecting the microorganism and cultivation conditions.

## 1. Introduction

The oils obtained from crops, especially palm oil, have been used as an inexpensive source for many useful components (like replacement of animal fats). It functions as a natural preservative in processed foods, is used as the foaming agent in shampoos and soaps, and used as a raw material for biofuels [1]. In the last two decades, the production of palm oil went up 241% and represented 37% of the global vegetable oil production in 2020 [2]. However, using these feedstocks is not ideal in the long term, as their production has severe consequences on biodiversity, and competes with other crops for land availability.

This impacts the food supply and hence, its sustainability is a major challenge due to the food crisis worldwide [3]. Furthermore, palm tree plantations are rapidly replacing the original tropical forests, and other original and traditional vegetation across numerous countries in Asia, South America, and Africa [4]. This is having a detrimental impact on the local ecosystem, as deforestation and climate change [5,6].

Despite the implementation of various responsible actions, including the opposition to deforestation driven by the RSPO (Roundtable on Sustainable Palm Oil), the utilization of palm oil remains a topic of considerable contention [7]. The use of palm oil (and other tropical oils) is growing annually by 4%, which cannot be provided by RSPO (without deforestation). Based on the Impact Report 2024 published by RSPO, only 8.1% of the palm oil production has RSPO-Certified [8]. To that end, implementing a sustainable alternative to fatty acids and oils derived from plant-based oils is urgent and of utmost interest. Oleaginous yeasts have a strong potential to produce sustainable alternatives to oils derived from plants due to their fast growth, high lipid content, and volumetric productivity [9]. Microbial oils produced by these oleaginous yeasts have been gaining significant attention as, third-generation biodiesel feedstocks, and a source for healthy oils/fatty acids such as PUFA’s (polyunsaturated fatty acids). These yeasts can store lipids (primarily as TAGs) ranging from 20% up to 80% of their cell mass under carefully selected culture conditions [10]. Under nitrogen limitation, fatty acid composition comprises myristic (C14:0, 0.3-34%), palmitic (C16:0, 3-37%), palmitoleic (C16:1, 0.4-9%), stearic (C18:0, 2-66%), oleic (C18:1, 36-57%), linoleic (C18:2, 2.1-24%), linolenic (C18:3, 1- 3%) acids [11]. Their versatile composition facilitates the use of oils in animal feed, food additives and ingredients, cosmetics, pharmaceuticals, and the production of biofuels, as they have a composition comparable to that of vegetable oils [12–14].

Over the last decade, research on yeast lipid technology has accelerated with a major interest focus on oleaginous yeast species, *Yarrowia lipolytica*, and *Cutaneotrichosporon oleaginosus*, as well as cultivation parameters to boost lipid accumulation, low-cost substrates, fatty acid composition, and cultivation modes to accelerate the industrial implementation of microbial oils [1]. Under nitrogen-limiting conditions, the composition of produced fatty acids has been reported to be 25% C16:0, 10% C18:0, 57% C18:1, and 7% C18:2 by *C. oleaginosus*, and 15% C16:0, 13% C18:0, 51% C18:1, and 21% C18:2 by *Y. lipolytica*: this is comparable to the composition of plant-based oils [16–18].

The fatty acid composition defines the physiological properties of the oil, thereby determining its potential applications. For instance, a higher proportion of saturated fatty acids contributes to the oxidative stability of the oil, the ratio of saturated to unsaturated fatty acids affects the melting point and a higher content of MUFAs improves the thermal stability of the oil [11,19,20]. The microbial lipids produced by *C. oleaginosus* and *Y. lipolytica* predominantly comprise C16 to C18 fatty acids. These fatty acids, particularly oleic acid (C18:1), render this an appealing raw material for the production of biodiesel [19,21]. Furthermore, oleic acid (C18:1) rich oils have been identified as superior feedstocks for chemical modification, facilitating the production of surfactants, plasticizers, and asphalt additives [20]. Saturated fatty acids such as C18:0 and C16:0-rich oils have been used in cleaning and personal care products [22,23]. For this reason, the production of microbial oil with an optimal compositional profile for a specific application is a critical aspect for ensuring its effective utilization.

In this study, we systematically investigated the variation in the fatty acid composition of the produced oil by *C. oleaginosus* and *Y. lipolytica* mutants at different cultivation parameters. Principal Component Analysis (PCA) and Covariance Simultaneous Component Analysis (COVSCA) were used to illustrate and interpret how correlation patterns vary across conditions (cultivation temperature, genetic interventions) and yeast species [24,25]. Following a thorough evaluation of the outcomes, the conditions were reported to obtain a specific fatty acid composition of the oil produced by *C. oleaginosus* and *Y. lipolytica*.

## 2. Materials and Methods

### 2.1. Strains and experimental conditions in literature

*Cutaneotrichosporon oleaginosus ATCC 20509, Yarrowia lipolytica CBS8108*, and transformants listed in Table 1 were used for cultivation. Wild-type (WT) and built *C. oleaginosus* and *Y. lipolytica* transformants were cultivated into minimal media consisting of glycerol as carbon source and urea as nitrogen source set ratios of C/N (g/g) 30, 120, 140, 175, 200, 240, or 300 [18,26–28]. Cultures were incubated at 15 °C, 30 °C, or 35 °C, at 250 rpm for 96 h (*C. oleaginosus* strains) and 120h (*Y. lipolytica* strains) in a shaking incubator. Cells were harvested at the end of incubation and centrifuged at 1780 g, 4 °C for 15 min. All experiments were performed in triplicates.

**Table 1.**
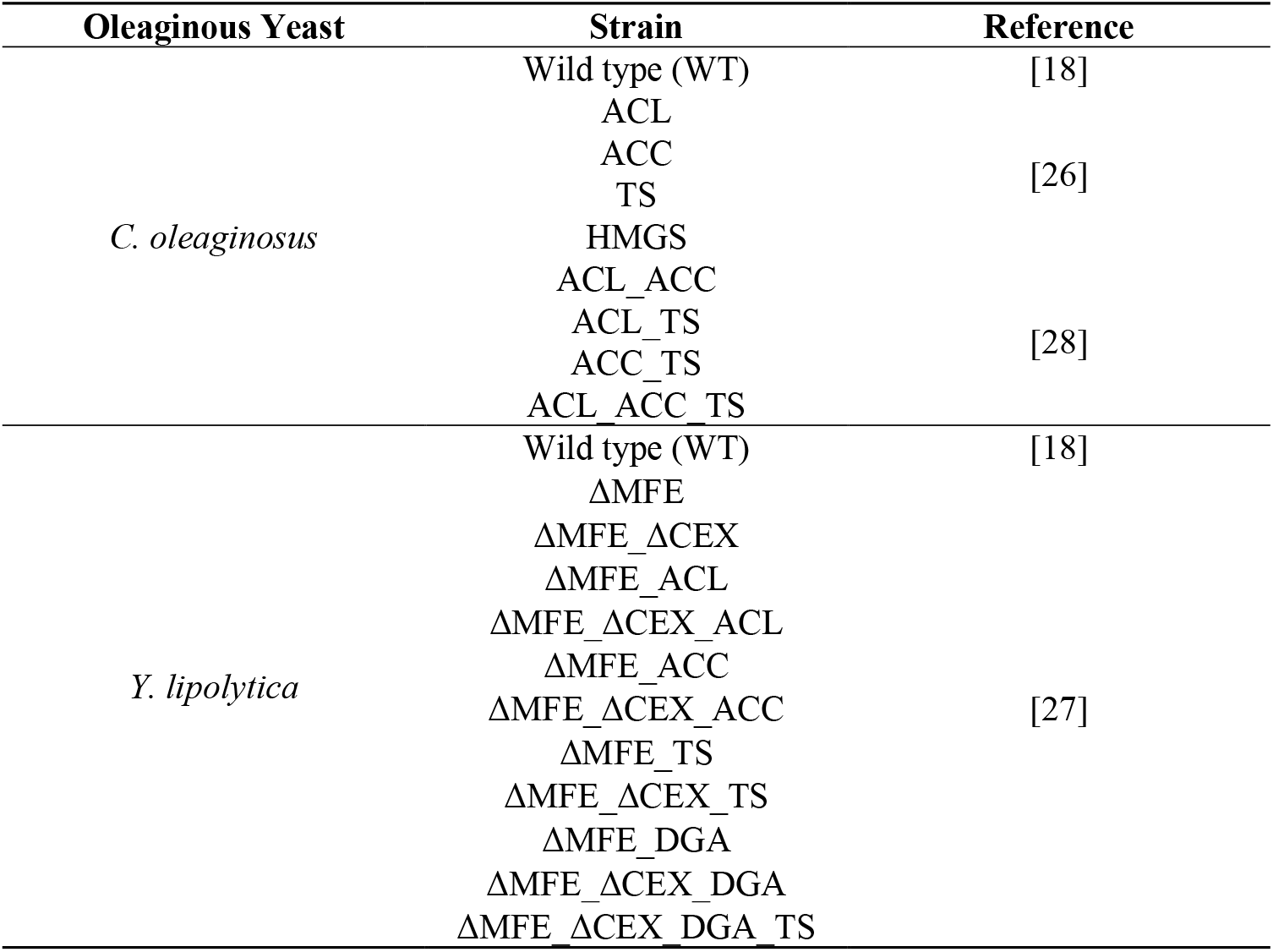
List of oleaginous yeast strains used in cultivation experiments.

### 2.2. Fatty acid composition data

The fatty acid composition of yeast microbial oil data was determined and acquired in the previous studies [18,26–28]. The total fatty acids were determined quantitatively with a gas chromatograph (GC), 7830B GC systems (Aligent, Santa Clara, CA, The US) equipped with a Supelco Nukol^TM^ 25,357 column (30 m x 530 µm x 1.0 µm; Sigma-Aldrich, St. Louis, MO, The US), hydrogen as a carrier gas. Samples were initially prepared as described by Duman-Özdamar et al., 2022 [18]. Chloroform was evaporated under nitrogen gas and lipid in the tubes was dissolved in hexane before GC analysis.

### 2.3. Statistical methods

Principal Component Analysis (PCA) [29,30] was carried out on the fatty acid profiles.

Covariance Simultaneous Component Analysis (COVSCA) [24] was performed to investigate the differences between the matrices of correlations obtained from the data matrix of *C. oleaginosus* and *Y. lipolytica*. For each microorganism, the full set of strains was considered, 9 and 12 for *C. oleaginosus* and *Y. lipolytica* respectively. For each strain matrices with the correlations between fatty acid types (9 variables) were built and differences were explored by fitting a COVSCA model considering 3 prototype matrices (rank 2).

### 2.4. Software and data

The statistics function prcomp within R version 4.0.2 [31,32]. The correlation biplots of the principal component scores, and the loading vectors and heatmaps were plotted through the R ggplot2 package [33].

Data used for PCA and COVSCA is available in ZENODO (https://doi.org/10.5281/zenodo.13958333), methods and code for COVSCA are available in

GitLab (https://github.com/esaccenti/covsca).

## 3. Results and Discussion

### 3.1. Fatty acid profile of oleaginous yeasts

The fatty acid composition of the microbial oils produced by *C. oleaginosus* and *Y. lipolytica* WT and mutants grown in minimal media with various C/N ratios at 30 °C were retrieved from our previous studies [18,26–28].

Oil from both oleaginous yeasts contained similar types of fatty acids including >1% C16:0, C16:1, C18:0, C18:1, C18:2, and C24:0, however, they also exhibited differences (Figure 1A). Microbial oil produced by *C. oleaginosus* contains a higher content of saturated fatty acids (35-44%, w/w) compared to the oil produced by *Y. lipolytica* (23-34%, w/w) when grown at 30 °C (Figure 1A, [ZENODO](https://wageningenur4-my.sharepoint.com/personal/efsun_duman_wur_nl/Documents/Tailor-made%20microbial%20oils/Papers/ShortCommunication_Intro_discussion/10.5281/zenodo.13958333)). The wild-type *Y. lipolytica* produced a higher content of PUFAs, with 20% (w/w) linoleic acid (C18:2), compared to the *C. oleaginosus* oil, which contains approximately 8% (w/w) C18:2 ([ZENODO](https://wageningenur4-my.sharepoint.com/personal/efsun_duman_wur_nl/Documents/Tailor-made%20microbial%20oils/Papers/ShortCommunication_Intro_discussion/10.5281/zenodo.13958333), Supplementary_file_A, Figure 5). On the other hand, the *C. oleaginosus* oil contains a higher content of MUFAs including 50% (w/w) oleic acid (C18:1) and 1% (w/w) palmitoleic acid (C16:1), whereas the *Y. lipolytica* oil comprises 36% (w/w) C18:1 and 9% (w/w) C16:1. Among tested *Y. lipolytica* mutants, we observed that ΔMFE_ΔCEX_DGA and ΔMFE_ΔCEX_DGA_TS presented a similar fatty acid composition to the oil produced by TS_ACC mutant of *C. oleaginosus*.

**Figure 1.**
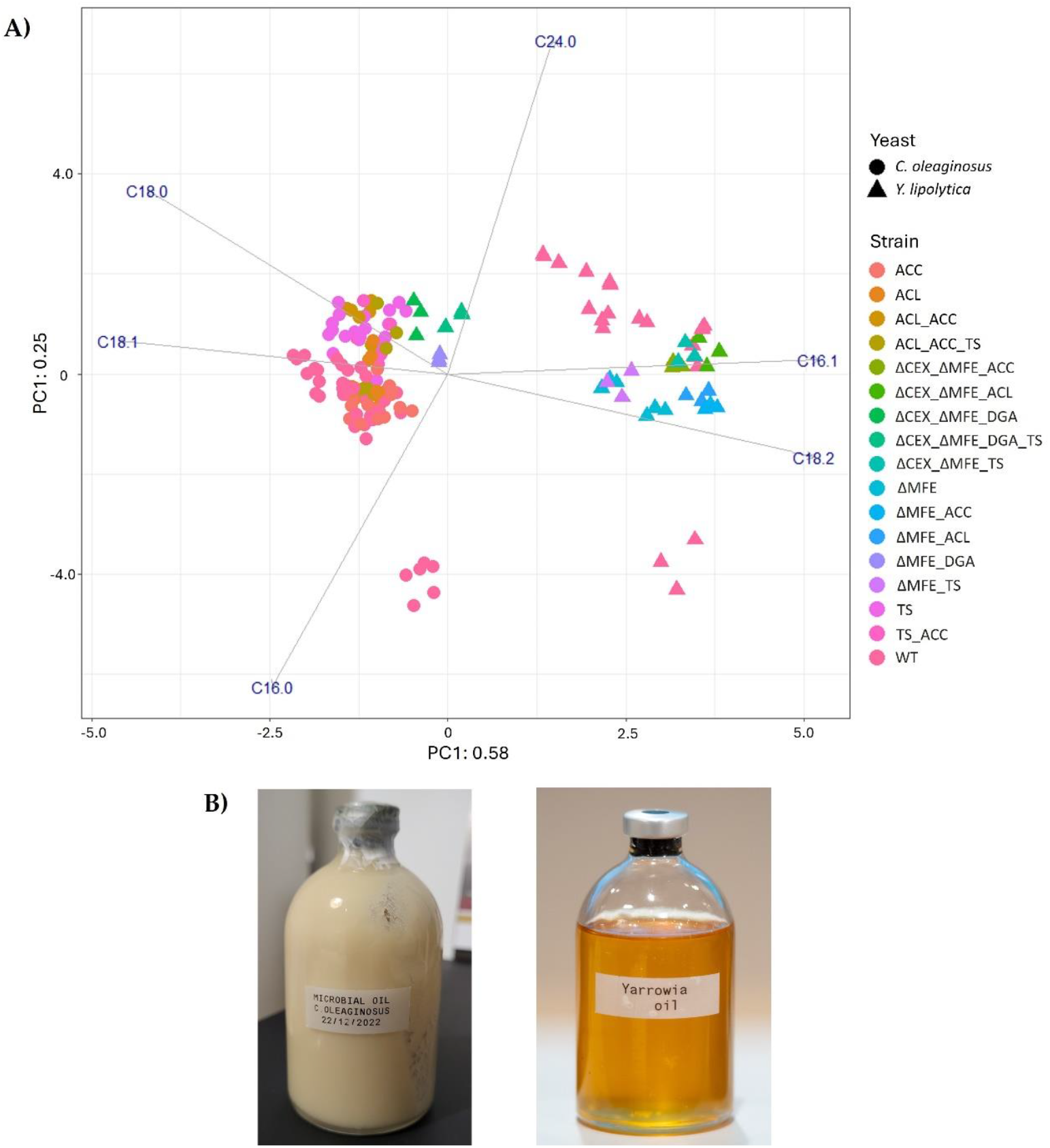
A) PCA on the fatty acid profile of the oil obtained from *C. oleaginous* and *Y. lipolytica*. In addition to the WT, different strains were included in the analysis. Variance explained by each component is given in the labels. **B)** Microbial oil obtained from *C. oleaginosus* (solid fat at room temperature) and *Y. lipolytica* (liquid oil at room temperature).

The observed variation in the ratio of saturated and unsaturated fatty acids resulted in oil with different melting points at room temperature. The extracted lipids from *C. oleaginosus* were solid at room temperature (fat), like palm oil (Figure 1B) [1]. Conversely, *Y. lipolytica* produced lipids that were liquid at room temperature (oil) (Figure 1B).

### 3.2. (Dis)similarities of the fatty acid composition produced by different strains

In addition to the oleaginous yeast-specific differences in the fatty acid profile, some of the applied genetic interventions and manipulated environmental factors steered the fatty acid composition. To investigate further the variation of fatty acid profiles among strains of oleaginous yeasts and to facilitate the appropriate strain selection, Covariance Simultaneous Component Analysis (COVSCA) was used to model the correlation matrices obtained for the lipids data of *C. oleaginosus* and *Y. lipolytica* were investigated.

The 3D plot in the COVSCA space (Figure 2) depicts the first three components of the model. Each point represents a strain. The axes (component space) show the dimensions where covariance is maximized, indicating that the closer the two points are, the more similar they are in terms of their correlation structure. For *C. oleaginosus*, wild-type and HMGS are clustered together, indicating the similarity in the fatty acid composition of the oil produced by these strains (Figure 2A). Although the fatty acid profiles of ACL, ACC, and TS were similar, the double and triple mutants ACL_TS and ACL_ACC_TS strains exhibited distinct fatty acid compositions compared to other strains. On the other hand, the fatty acid composition of *Y. lipolytica* strains exhibited greater divergence except for ΔMFE_ΔCEX_ACL, ΔMFE_ΔCEX_ACC, and ΔMFE_ΔCEX_TS, which represented similarities (Figure 2B). Furthermore, ΔMFE_ΔCEX_DGA_TS showed strong variation along all axes, indicating that combining multiple genetic modifications resulted in a significant alteration in the fatty acid profile. In all, COVSCA models and plots demonstrate the possibility of creating a pallet of strains expressing a wide range of fatty acid profiles.

**Figure 2.**
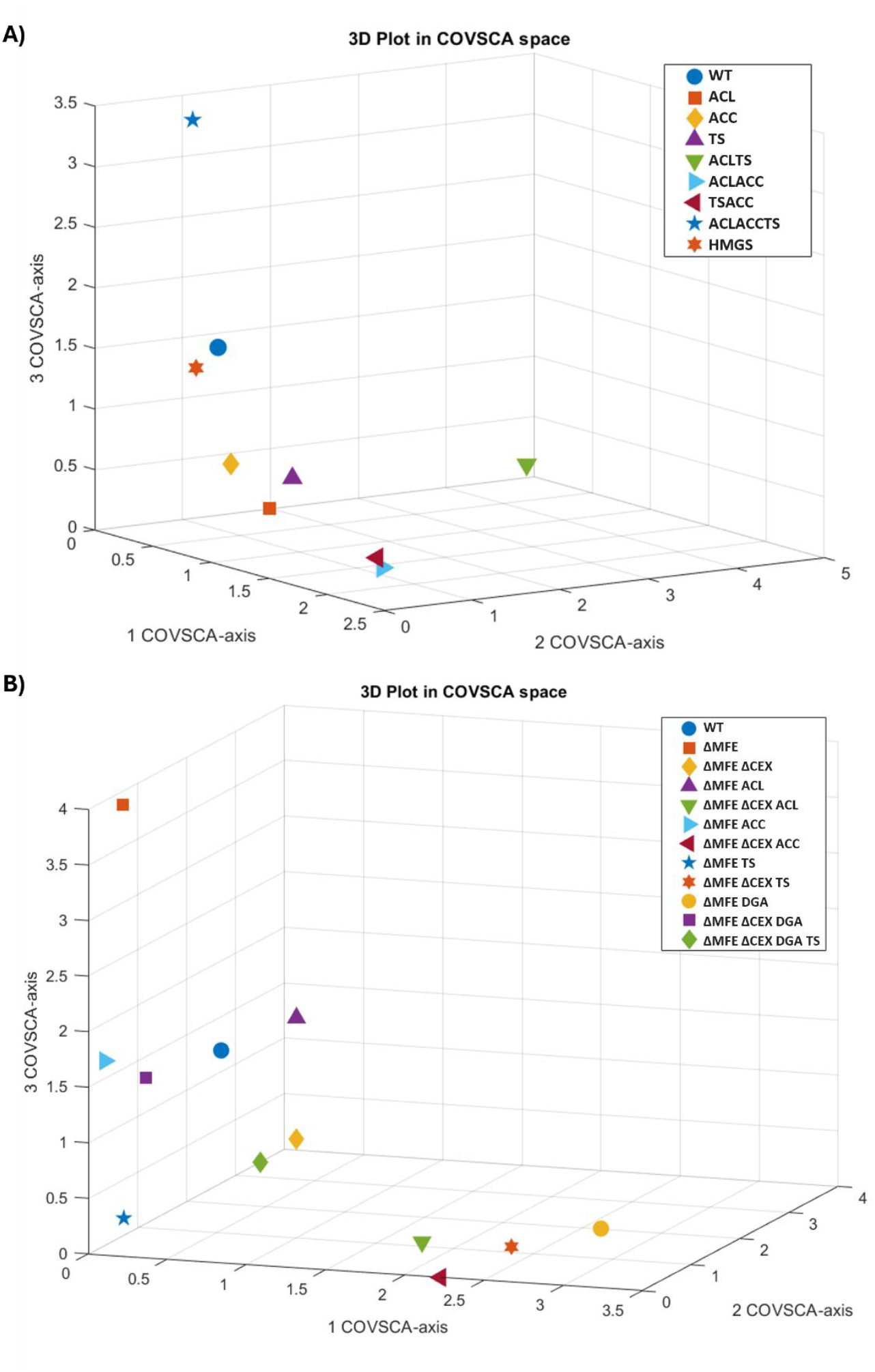
Plots in COVSCA Space representing the (dis)similarities of the fatty acid composition obtained by incubating various strains of **A)** *C. oleaginosus* and **B)** *Y. lipolytica* at different temperatures and C/N ratio medium. COVSCA models of *C. oleaginosus* and *Y. lipolytica* strains had 75.10% and 57.36% goodness of fit respectively.

### 3.3. Altered cultivation parameters and genetic modifications steered fatty acid composition

A comprehensive comparison of all tested conditions (temperature, C/N ratio, and medium supplements) and applied genetic modifications was conducted for each fatty acid component (Figure 3). We observed that varying the cultivation temperature steered the fatty acid composition (Figure 3). For both oleaginous yeasts, when the growth temperature was increased to 35 °C, they produced a higher content of saturated fatty acids (C18:0, C20:0, C24:0). Bellou et al. (2016) reported that temperature affects the activity of fatty acid desaturase enzymes therefore the fatty acid composition of produced microbial oil is altered [34].

**Figure 3.**
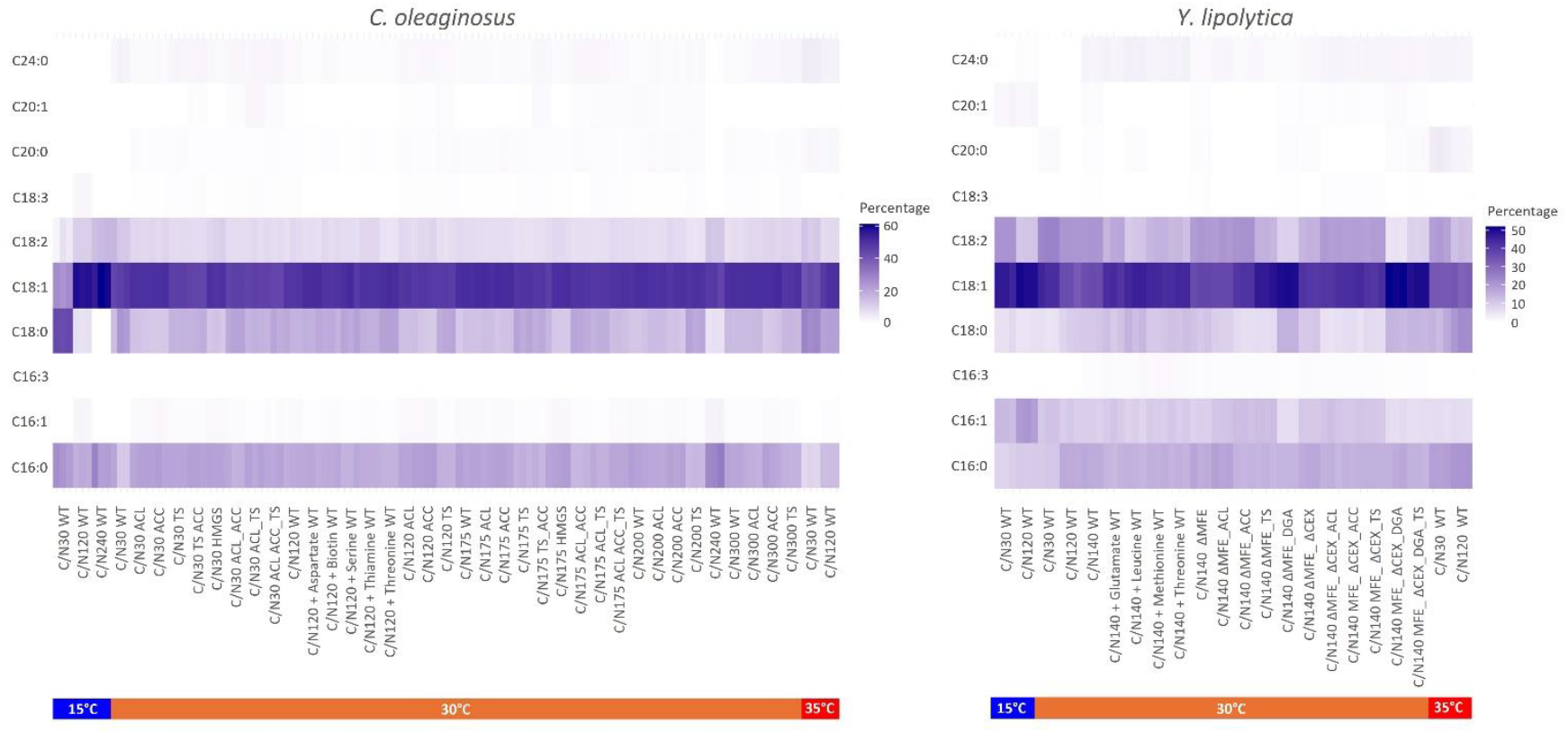
Heatmap of the fatty acid profile of *C. oleaginosus* and *Y. lipolytica*. For each experimental condition triplicates were used in the plots. The temperatures are included as bars below the sample names represented by colors.

Moreover, an increased C18:0 fraction in total fatty acid was observed when *TS* was overexpressed in *C. oleaginosus* (TS, ACL_TS, ACC_TS, ACL_ACC_TS) compared to other strains (Figure 3). On the other hand, overexpression of *TS* in *Y. lipolytica* led to the reduction of the C18:0 fraction. Enzyme inhibitors such as specific fatty acids can be added to the broth to alter the lipid fatty acid profile [35,36]. This strategy was considered when attempting to increase the stearic acid (C18:0) content for the production of a cocoa butter equivalent (CBE) with *C. oleaginosus*, but they increase process costs and some are toxic [37,38]. To encounter the increased cocoa butter prices, in addition to CBE, cocoa butter replacers (CBR) are developed as alternatives. CBRs are produced by hardening the oils derived from sunflower or rapeseed via partial hydrogenation and therefore comprise a high amount of trans-fat [39]. In this regard, by altering the temperature and selecting the appropriate strain, the oil replacement for tropical (hard) fats, such as cocoa butter, palm, and coconut fat can be obtained via *C. oleaginosus*.

Decreasing the temperature to 15 °C led to the production of a higher content of unsaturated fatty acids (C18:1, C18:2). Gao et al. (2020), tested 28 °C and 38 °C for *Y. lipolytica* and reported a similar shift in the fatty acid composition of the produced oil [40]. Among all tested strains and cultivation conditions, the highest C18:1 composition (∼60%, w/w) was obtained with wild-type *C. oleaginosus* incubated at 15 °C in C/N 120 medium ([ZENODO](https://doi.org/10.5281/zenodo.13958334), Supplementary File A, Figure 4). For *Y. lipolytica*, C18:1 composition reached up to ∼50% (w/w) when ΔMFE_DGA, ΔMFE_ΔCEX_DGA, and ΔMFE_ΔCEX_DGA_TS were cultivated at 30 °C in C/N140 medium and the wild-type was incubated at 15 °C ([ZENODO](https://doi.org/10.5281/zenodo.13958334), Supplementary File A, Figure 9). High relative oleic acid concentrations (∼50% (w/w)) were observed with oleaginous yeasts that accumulate more than 50% lipid content. Tai et al. (2013) reported higher C18:1 composition when *DGA1* was overexpressed and related this situation with rapid lipid production in that strain [41].

Furthermore, the supplementation of amino acids to the medium with wild-type strains did not result in a change in the ratio of saturated and unsaturated fatty acids. However, this led to a shift in the profile of PUFAs towards MUFAs, with a notable increase in the fraction of C18:1 (Figure 3). Tsakraklides et al. (2018) obtained lipid accumulation with a C18:1 content exceeding 90% in *Y. lipolytica* by replacing the native Δ9 fatty acid desaturase and glycerol-3-phosphate acyltransferase with heterologous versions [42,43]. Additionally, the Δ12 fatty acid desaturase was deleted and a fatty acid elongase was expressed. The combination of these approaches can be beneficial in increasing the content of oleic acid in both oleaginous yeasts. Furthermore, by decreasing the temperature, the ratio of unsaturated fatty acids can be increased in *C. oleaginosus* thereby potentially enabling the production of oil with a similar composition to that of *Y. lipolytica*. However, it should be noted that reduced temperature may have an impact on the overall yield of microbial oil production [18].

In this work, we investigated the fatty acid composition of the microbial oil by using glycerol as a carbon source and urea as a nitrogen source. Awad et al. (2019), reported that using different carbon and nitrogen sources influenced the saturation level of the fatty acids produced by *C. oleaginosus* [44]. The supplementation of ammonia in the media resulted in a notable shift in the fatty acid profile of the produced oil compared to the profiles resulting from the supplementation of urea, yeast extract, and peptone (organic nitrogen sources). Using ammonia increased the proportion of C18:2 and decreased the proportion of C16:0 and C18:0. Additionally, Awad et al. (2019) observed comparable fatty acid profiles when glucose, mannose, maltose, and lactose were utilized as carbon sources, similar to the findings observed with glycerol [44]. Conversely, sorbitol and arabinose reduced C18:0 and C18:1, with an increase in C18:2. For *Y. lipolytica*, dextrose was found to enhance the content of saturated fatty acids (C16:0 and C18:0) [45]. This represents that specific fatty acid profiles associated with desired product applications can be boosted by carefully selecting media components.

## 4. Conclusions

The systematic evaluation of all tested conditions (temperature, C/N ratio, and medium supplements) and applied genetic modifications to improve the lipid accumulation of *C. oleaginosus* and *Y. lipolytica* revealed the variation that occurred in fatty acid composition. These findings suggest that combining optimized media composition, and cultivation conditions with genetically engineered strains enables achieving unique combinations of both saturated and unsaturated fatty acids in significant quantities. This will facilitate the utilization of microbial oil from *C. oleaginosus* and *Y. lipolytica* as a substitute for plant-based oils and fats for versatile industrial applications while still maintaining a high lipid content.

### Supplementary Materials

Supplementary files are available at ZENODO doi: https://zenodo.org/records/13958334.

## Abbreviations

The following abbreviations are used in this manuscript:

C/N: Carbon to nitrogen
PCA: Principal Component Analysis
COVSCA: Covariance Simultaneous Component Analysis

## Author Contributions

Conceptualization, ZEDÖ, JH, VAPMS, MSD; methodology, ZEDÖ, ES, MSD; formal analysis, ZEDÖ, ES, MSD; investigation, ZEDÖ, JH, MSD; data curation, ZEDÖ, ES, MSD; writing—original draft preparation, ZEDÖ; writing—review and editing, ZEDÖ, ES, VAPMS, JH, MSD; visualization, ZEDÖ, ES, MSD; supervision, VAPMdS, JH, MSD; project administration, VAPMdS, JH; funding acquisition, VAPMdS, JH, MSD. All authors have read and agreed to the published version of the manuscript.

## Funding

This research was financed by the Dutch Ministry of Agriculture through the “TKI-toeslag” project LWV19221 “Tailor-made microbial oils and fatty acids”.

## Data Availability Statement

Supplementary files are available at ZENODO doi:

https://zenodo.org/records/13958334.

## Conflicts of Interest

JH has interests in NoPalm Ingredients BV and VAPMdS has interests in LifeGlimmer GmbH.

